# Assessing the role of BNST GABA neurons in backward conditioned suppression

**DOI:** 10.1101/2025.01.01.631006

**Authors:** Annie Ly, Hayden Hotchkiss, Emily D. Prévost, Julianne Pelletier, Melissa Deming, Luma Murib, David H. Root

**Author notes:** Correspondence: David H. Root, Ph.D., Boettcher Investigator, Department of Psychology and Neuroscience, University of Colorado Boulder, 2860 Wilderness Place Boulder, CO 80301 USA.

## Abstract

Conditioned suppression is a useful paradigm for measuring learned avoidance. In most conditioned suppression studies, forward conditioning is used where a cue predicts an aversive stimulus. However, backward conditioning, in which an aversive stimulus predicts a cue, provides unique insights into learned avoidance due to its influence on both conditioned excitation and inhibition. We trained mice to consume sucrose in context A, associated an aversive stimulus in context B to few or many forward or backwards paired cues (CS+), and then tested for conditioned suppression in context A in response to the CS+. We found that few or many forward CS+ and few backward CS+ produced conditioned suppression, but many backwards cues did not. Administration of diazepam, a positive allosteric modulator of the GABA-A receptor, prevented conditioned suppression to the backward CS+ but not to the forward CS+. Furthermore, freezing behavior was observed in response to the forward CS+ but not the backward CS+, and diazepam had no effect on freezing or locomotion. We next examined BNST GABA neurons for potential sensitivity to backwards cues and conditioned suppression. VGaT BNST signaling increased in response to sucrose licks during the backward CS+ but not to licks outside the CS+ and not to the backward CS+ onset or offset. Using designer receptors, we found that BNST VGaT neuron activation, but not its inhibition, prevented backward conditioned suppression expression. We conclude that backward conditioned suppression is dependent on both positive allosteric modulation of GABA on GABA-A receptors by diazepam and BNST GABA neurons.

## INTRODUCTION

Conditioned suppression is a psychology-based task to understand how learned cues interrupt ongoing reward-seeking behavior. In conditioned suppression studies, the presentation of a cue that was previously paired with an aversive outcome is sufficient to suppress a reward response, whether that response is operant (Estes & Skinner, 1941) or consummatory (Leaf & Muller, 1965) by design.

Traditionally, conditioned suppression studies use forward conditioning pairings in which a cue (CS+) predicts a forthcoming unconditioned aversive stimulus (US) (Arico et al., 2017; Bercum et al., 2021; Bouchekioua et al., 2022; Bouton & Bolles, 1979; Greiner et al., 2019; Piantadosi et al., 2020). Forward conditioning effectively suppresses consummatory behavior during CS+ (Leaf & Muller, 1965). Regardless of the number of forward CS+ pairings, a forward CS+ remains a conditioned excitor. In the case of conditioned suppression, the forward paired CS+ excites the predicted conditioned response associated with the US, typically freezing behavior and reduced reward-related responding (Leaf & Muller, 1965). Backward conditioning, where an aversive stimulus predicts a forthcoming CS+, has similarities and differences to forward conditioning. Like forward conditioning, a few backward pairings result in the CS+ becoming an excitatory stimulus (evoking a conditioned response). However, many backward pairings results in the CS+ becoming an inhibitory stimulus (not evoking a conditioned response) (Ayres et al., 1987; Barnet & Miller, 1996; Chang et al., 2003; Christianson et al., 2008; Cole & Miller, 1999; Heth, 1976; Heth & Rescorla, 1973; Maier et al., 1976; Moscovitch & LoLordo, 1968; Spetch et al., 1981). Rescorla defined a conditioned inhibitor as a CS+ that inhibits the expected conditioned response due to the learned association with the absence of the US (Christianson et al., 2012; Rescorla, 1969). Others have used this definition of conditioned inhibition to investigate the learning and memory of “safety” (Ng et al., 2024; Sangha et al., 2013; Sangha et al., 2014). Wagner offered an explanation for the unique shift from conditioned excitation to conditioned inhibition feature of backward conditioning with the sometimes-opponent process theory, which posits a trial-based acquisition of the conditioned response (Wagner, 1981). Others have utilized this feature of backward conditioning to create a “safety signal” with many backward CS+ pairings (Christianson et al., 2008). While conditioned response characteristics about forward and backward conditioning are well studied, how the history of backwards conditioning may differ from forward conditioning with respect to conditioned suppression is unknown.

Freezing is the most readily observed behavior in response to forward conditioning of an aversive stimulus, while locomotor activity is observed in response to backward conditioning of an aversive stimulus (Bouton & Bolles, 1980). Indeed, greater time spent freezing appears to correlate to more conditioned suppression (Bouton & Bolles, 1980; Mast et al., 1982). Despite involving similar behaviors, there is support for the idea that conditioned freezing and conditioned suppression are mutually exclusive behaviors that are mediated by different neural circuits (Amorapanth et al., 1999; Blanchard et al., 2001; Killcross et al., 1997; McDannald, 2010; McDannald & Galarce, 2011).

The bed nucleus of the stria terminalis (BNST) is an extended amygdala structure that is cellularly heterogeneous but predominantly GABAergic (Bota et al., 2012; Siletti et al., 2022; Welch et al., 2019). In comparisons between few pairings of backward and forward conditioning, the BNST is strongly implicated in the processing and conditioned responding to ambiguous threats such as few backward paired CS+ (Goode et al., 2020; Goode et al., 2019; Ressler et al., 2020). Here, we compared conditioned suppression between a few and many pairings of forward and backward CS+ that predicted an aversive stimulus and investigated the role of BNST GABA neurons in conditioned suppression evoked by backwards CS+. Consistent with the trial-based model of the sometimes-opponent process theory, the forward CS+ suppressed sucrose consumption regardless of the number of pairings, and a few pairings of backward CS+ but not many pairings of the backward CS+ suppressed sucrose consumption. Diazepam selectively prevented conditioned suppression to few pairings of the backward CS+ and not to the forward CS+. Fiber photometry GCaMP recordings showed that BNST GABA neurons signaled sucrose consumption during few pairings of the backward CS+ but not to sucrose consumption without the backward CS+, or the onset or offset of the backward CS+ itself. Designer receptor activation of BNST GABA neurons, but not designer receptor inhibition of BNST GABA neurons, abolished backward conditioned suppression. The results described here suggest that, while both forward and backward conditioning can produce conditioned suppression, 1) they are distinguishable by their dependency on diazepam and the behaviors that occur during cue presentation during conditioned suppression; and 2) backward conditioned suppression is dependent on both positive allosteric modulation of GABA on GABA-A receptors by diazepam and the activation of BNST GABA neurons.

## MATERIALS AND METHODS

### Animals

Wildtype mice (4-5 months old) were used for the comparison between few and many forward and backward pairings on conditioned suppression (N=33; 13 female, 20 male), as well as for the effect of diazepam (N=41, 20 female, 21 male). VGaT-IRES::Cre knock-in mice (4-5 months old, Slc32a1tm2(cre)Lowl/J; Stock #016962) were purchased from The Jackson Laboratory (Bar Harbor, ME) and bred at the University of Colorado Boulder. VGaT-IRES::Cre mice were used for fiber photometry (N=8, 4 female, 4 male) and designer receptors exclusively activated by designer drugs (DREADDs) manipulation (N=22). Mice were group-housed by sex (4-5 mice/cage) under a reversed 12hr:12hr light/dark cycle (lights on at 10pm) with access to water ad libitum. All mice were weighed daily and fed to maintain 85% of their body weight. Food-restricted mice were fed after sucrose access. All experiments were performed during the dark phase of the light cycle. The experiments described were conducted in accordance with the regulations by the National Institutes of Health Guide for the Care and Use of Laboratory Animals and approved by the Institutional Animal Care and Use Committee at the University of Colorado Boulder.

### Surgery

During surgery, VGaT-IRES::Cre mice were continuously anesthetized with 1-2% isoflurane gas while secured in the stereotactic instrument. Mice were injected with AAV1-hSyn-FLEX-GCaMP6m (Addgene) unilaterally into the BNST (5 × 1012 titer, 350 nL volume; 100 nl/min rate; +0.3 mm anteroposterior, +0.6 mm mediolateral, −4.1 mm dorsoventral coordinates from bregma) using an UltraMicroPump, Nanofil syringes, and 35-gauge needles (Micro4; World Precision Instruments, Sarasota, FL). For DREADD experiments, VGaT-IRES::Cre mice were bilaterally injected with either AAV8-hSyn-DIO-mCherry, AAV8-hSyn-DIO-hM4Di-mCherry, or AAV8-hSyn-DIO-hM3Dq-mCherry using the same coordinates as previously described. Syringes were left in place for 10 min following injections before being slowly withdrawn. For fiber photometry, after the syringe was withdrawn, an optic fiber (400μm core diameter, 0.66 NA, Doric Lenses) was implanted slightly dorsal to the BNST (+0.3 mm anteroposterior, ±0.6 mm mediolateral, −3.9 mm dorsoventral coordinates from bregma). All implants were secured with skull screws and dental cement. Mice were given 3 days of postoperative care with daily carprofen (5 mg/kg, I.P.) and allowed 3-4 weeks of recovery before experimentation.

### Histology

At the conclusion of fiber photometry experiments, VGaT-IRES::Cre mice were perfused transcardially with 0.1M phosphate buffer followed by 4% (w/v) paraformaldehyde in 0.1M phosphate buffer, pH 7.3. Brains were extracted and cryoprotected in 18% sucrose solution in 0.1M phosphate buffer at 4°C overnight. Brains were cryo-sectioned to obtain coronal slices with BNST (30 μm). These coronal brain slices were mounted onto gelatin-coated slides and imaged for GFP or mCherry fluorescent expression on a Zeiss widefield Axioscope at 5X magnification.

### Pavlovian Conditioning

In behavior chambers (Med-Associates) outfitted with a spout for a 250ml sipper bottle (Allentown, LLC) filled with 8% sucrose solution, mice were given 10-minute access over the course of 10 consecutive days. After 10 sessions of free access to sucrose, mice were placed in a wheel-turn chamber (Med-Associates) for Pavlovian conditioning acquisition. The wheels were locked and therefore inoperable to the mice. Using surgical tape, the tails of the mice were taped down to a Plexiglas rod and affixed with copper electrodes and electrode cream. Mice were either exposed to 12 forward CS+ pairings (12 FW), 12 backward CS+ pairings (12 BW), 96 forward CS+ pairings (96 FW), 96 backward CS+ pairings (96 BW), or no shocks as control (CTL). The 12 pairings were determined based on prior published work deeming sufficient conditioned suppression with this number of forward and backward pairings (Goode et al., 2020; Goode et al., 2019; Ressler et al., 2020). The 96 pairings were determined based on published work finding that this number of pairings was sufficient to turn backward conditioning into a conditioned inhibitor (Christianson et al., 2008; Cole & Miller, 1999; Heth, 1976; Maier et al., 1976; Moscovitch & LoLordo, 1968). The CS+ for all experiments was an auditory tone (80dB, 2KHz, 10 sec, random intertrial interval 60 sec), which was paired with the US (tail shock, 0.3 mA, 2 sec duration). The forward CS+ offset equaled the US onset. The backward CS+ onset equaled the US offset. Previous work has demonstrated that a 0 sec trace interval produced conditioned suppression (Marlin, 1981). After conditioning, mice were returned to their home cage. The following day, they were placed back into the behavior chambers with free access to sucrose. 15 CS+ presentations (80dB, 2KHz, 10 sec, random intertrial interval with an average of 60 sec) were played in the behavior chamber.

For diazepam experiments, mice were injected with 0.5mg/kg of diazepam 30 minutes before being exposed to the CS+ in the behavior chambers. In other studies, mice that were injected with 0.5mg/kg diazepam were able to maintain rotarod performance, but mice injected with 1.0mg/kg or higher doses of diazepam experienced greater diminishment in rotarod performance (Rosland et al., 1987). Therefore, we proceeded with a diazepam dose of 0.5 mg/kg.

### Calcium Fiber Photometry Recordings

GCaMP6m was excited at two wavelengths (465nm and 405nm isosbestic control) with amplitude-modulated signals from two light-emitting diodes reflected off dichroic mirrors and then coupled into an optic fiber (McGovern et al., 2024; McGovern et al., 2021). The GCaMP signal and the isosbestic control signal were returned through the same optic fiber and acquired using a femtowatt photoreceiver (Newport, Irvine, CA), digitized at 1kHz, and then recorded by a real-time signal processor (Tucker Davis Technologies). For analysis of calcium fiber photometry recordings, custom-written MATLAB scripts were used and are available at www.root-lab.org/code. The isosbestic signal (405nm) and the GCaMP signal (465nm) were downsampled (10x), and peri-event time histograms were created surrounding each event. For each trial, data were detrended by regressing the 405nm signal on the 465nm signal. The generated linear model was used to create a predicted 405nm signal that was subtracted from the 465nm signal to remove movement, photo-bleaching, and fiber bending artifacts (Barker et al., 2017). Baseline normalized maximum z-scores were taken from −5 to 0.01 seconds prior to event onset. A lick bout was defined as the occurrence of at least 2 licks. Onset of a lick bout was defined as the first lick with at least 3 sec of no licks prior, and the offset of a lick bout was defined as the last lick with a following interval of at least 400 msec. For the first lick during the CS+, baseline normalized z-scores were taken from −13 to −10 sec prior to cue onset. Intertrial licks or uncued licks are defined as all licks that did not occur during CS+.

### Statistical Analysis

Values are reported as mean ± standard error with standard error represented as bars or shaded areas. All statistical tests were performed with R (4.0.5). All results were subject to a two-way ANOVA analysis, where appropriate. In the case of evaluating change over time with session effect, an ANOVA was run on a linear mixed effects model with subject by session as the random effect. When there were no sex differences, analyses were pooled across sex thereafter. Post-hoc analysis was made by TukeyHSD.

## RESULTS

### Forward CS+ and few pairings of backward CS+ produced conditioned suppression

Wildtype mice were given 10-minute free access to 8% sucrose solution from sipper bottles in behavior chambers (Context A) for 10 consecutive days. There were neither sex differences nor group differences for total number of licks in the last 3 sessions of sucrose access (**Figure 1A and 1B**). After 10 sessions, mice were placed in a different chamber (Context B) that allowed for Pavlovian conditioning using tail shocks as an unconditioned stimulus. Mice either received 12 forward CS+ pairings (12 FW), 12 backward CS+ pairings (12 BW), 96 forward CS+ pairings (96 FW), 96 backward CS+ pairings (96 BW), or no shocks and no cues as a control (CTL). One day after Pavlovian conditioning, mice were placed back in the sucrose consumption chamber (Context A) with CS+ presentations to test for conditioned suppression (**Figure 1C**). A between-subjects ANOVA yielded a group effect in the total number of licks during the CS+ presentations [F(4,28) = 4.22, p = 0.008; 12 FW = 32.1±8.45, 96 BW = 36.1±14.4, 12 BW = 50.5±11.6, 96 BW = 108.1±25.8, CTL = 87.2±7.74] (**Figure 1D**).

**Figure 1.**
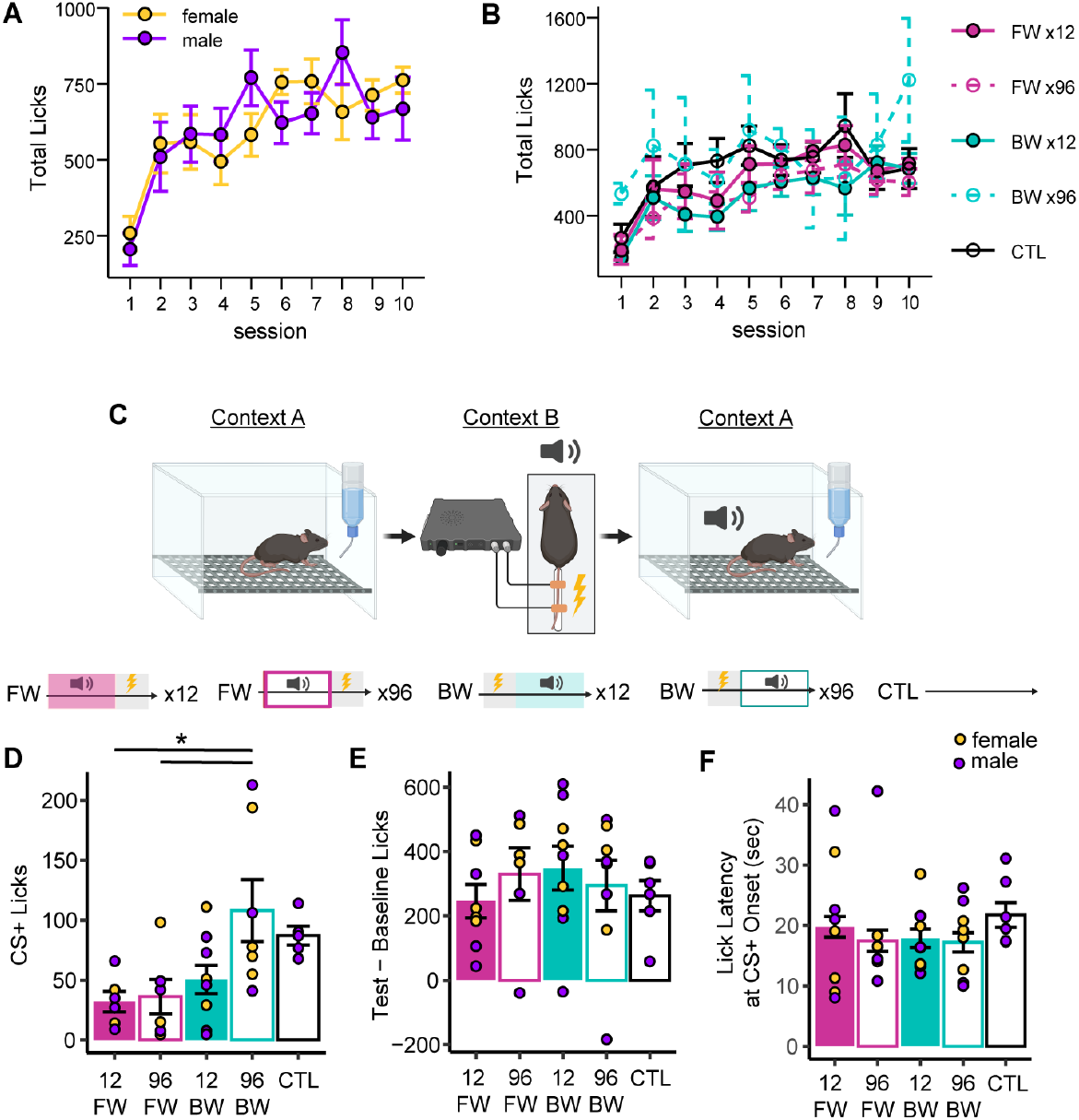
Forward and few backward paired cues cause conditioned suppression. (A) Total licks of sucrose between females and males before conditioning. (B) Total licks of sucrose between groups before conditioning. (C) Illustration of experimental timeline and groups. (D) Total licks during CS+ in Context A between groups. (E) Difference in total licks during the conditioned suppression test subtracted from baseline between groups. (F) Latency to first lick after CS+ offset. * p < 0.05

Posthoc Tukey tests showed that 96 BW mice had significantly higher licks during the CS+ than 12 FW mice [p = 0.02] and 96 FW mice [p = 0.02]. The difference between 12 and 96 backward pairings was near significance [p = 0.06]. Notably, there was no difference in cued licks between 12 BW and the FW groups, nor differences in the cued licks between 96 BW and CTL mice. There were no group differences in the change in licks from the day before conditioning (baseline) to total licks, including cued and uncued, during the conditioned suppression test day (**Figure 1E**). Furthermore, there were no group differences in the latency to lick following the CS+ offset (**Figure 1F**), indicating that CS+ presentations did not suppress licking outside of the cues.

### Diazepam prevented conditioned suppression to few pairings of backward CS+

Diazepam is a benzodiazepine that is a well-established treatment option for anxiety disorders (WHO Model List of Essential Medicines - 23rd list, 2023), and its mechanism of action is through the positive allosteric modulation of GABA to the GABA-A receptor (Calcaterra & Barrow, 2014). To understand the role of GABA signaling in conditioned suppression, wildtype mice were first trained to consume sucrose in context A. Afterwards, they underwent Pavlovian conditioning in context B. The following day, they were systemically injected with 0.5mg/kg diazepam (DIAZ) or vehicle solution (VEH) before being placed in context A while exposed to CS+ presentations and consuming sucrose reward (**Figure 2A**). Here, mice only had 12 forward CS+ pairings (12 FW) or 12 backward CS+ pairings (12 BW). The between-subjects ANOVA yielded a significant interaction between group and treatment [F(1, 37) = 10.5, p =0.002; FW VEH = 37.0±11.9, BW VEH = 89.1±17.9, FW DIAZ = 21.5±4.95, BW DIAZ = 182.0±20.9] (**Figure 2B**). Post-hoc analysis revealed that diazepam blocked conditioned suppression specifically in mice that received backwards conditioning (BW mice) [p = 0.001] and not in forwards conditioned mice (FW). There were no group differences in cued licks between FW and BW when treated with VEH, and diazepam had no effect on FW mice. To examine if nonlicking behaviors were altered differently between FW and BW groups, and the effect of diazepam of them, we examined freezing and locomotor behavior during CS+ presentations. We found an overall significant difference in freezing between FW and BW groups [F(1, 42) = 28.0, p < 0.0001; FW = 106.4±4.84, BW = 70.5±5.11] (**Figure 2C**). However, there was no diazepam treatment or interaction effect on freezing behavior. There was also a group difference in distance traveled during CS+ [F(1, 44) = 27.9, p < 0.0001; FW = 0.45±0.06, BW = 1.10±0.10], but there was neither a diazepam effect nor interaction of diazepam treatment with groups (**Figure 2D**). There were no group differences in total distance traveled (**Figure 2E**), average speed (**Figure 2F**), and time spent oriented towards the sucrose spout for the whole session (**Figure 2G**). Together, the positive allosteric modulation of GABA to the GABA-A receptor by way of diazepam treatment prevented the conditioned suppression effect to backward CS+ without altering forward CS+, locomotion, or freezing behavior.

**Figure 2.**
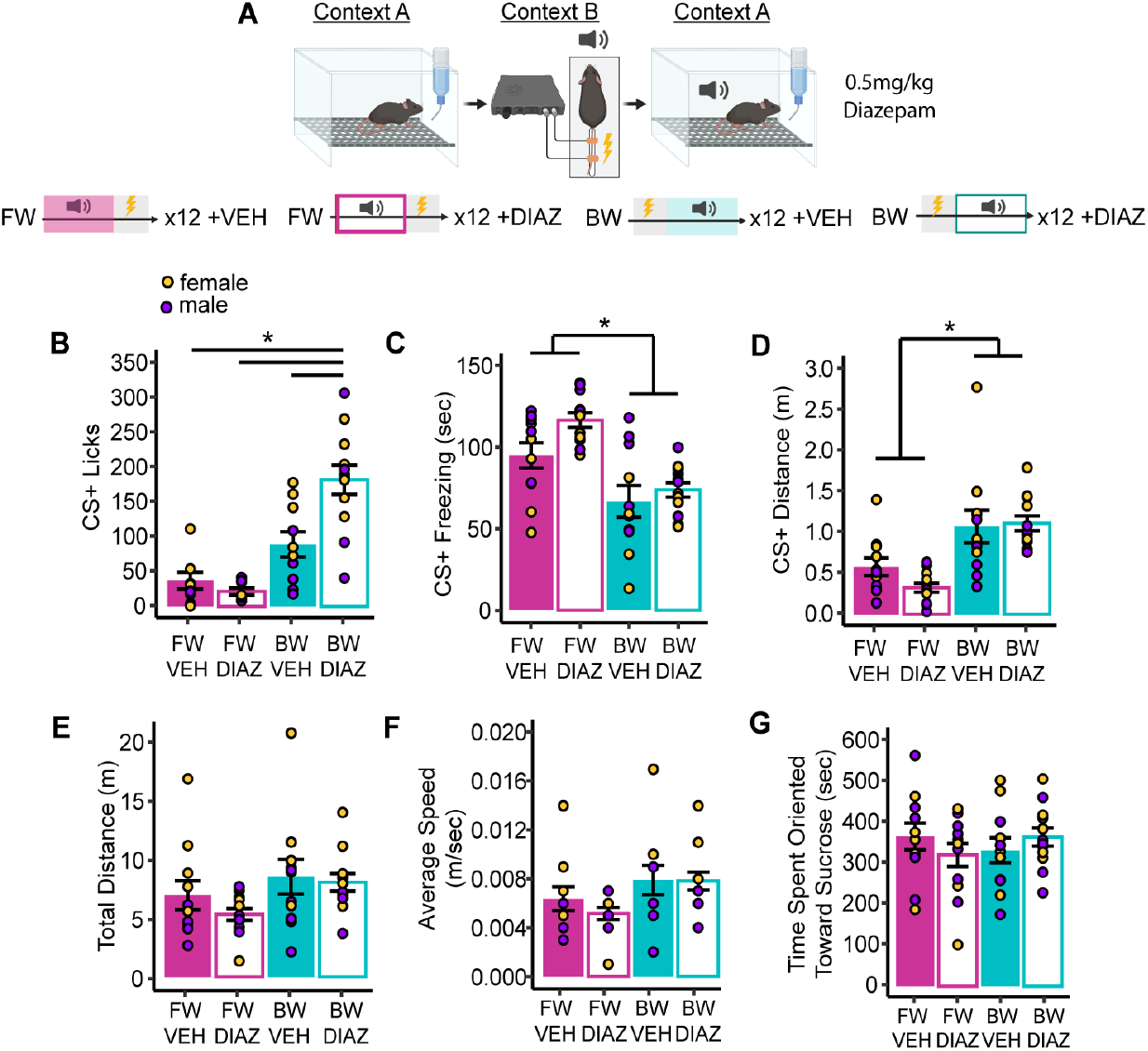
Backward conditioned suppression is blocked by diazepam. (A) Illustration of experimental design of diazepam I.P. treatment during CS+ the conditioned suppression test for groups that underwent 12 pairings of forward or backward CS+. (B) Total licks during CS+ the conditioned suppression test. (C) Freezing behavior during CS+ the conditioned suppression test. (D) Distance traveled during CS+ the conditioned suppression test. (E) Total distance traveled. (F) Average speed. (G) Time spent oriented toward the sucrose sipper bottle. * p < 0.05

### VGaT BNST neurons signal to sucrose licks during few paired backward CS+ but not to all licks

As a highly GABAergic brain region implicated in anxiety-like behaviors, we recorded the GCaMP neuronal activity of BNST neurons expressing the vesicular GABA transporter (VGaT) during few paired backward CS+ presentations in the conditioned suppression task (**Figure 3A and 3B**). We averaged traces between backward CS+ licks and uncued licks (**Figure 3C**), as well as traces to backward CS+ onset and offset (**Figure 3D**). There was a significant difference in the maximum peak of calcium-dependent signaling between cued licks, uncued licks, CS+ onset, and CS+ offset [F(3, 28) = 11.1, p <0.0001; CS+ licks = 4.17±0.88, uncued licks = 0.61±0.12, CS+ onset = 1.61±0.25, CS+ offset = 1.06±0.21] (**Figure 3E**). In post-hoc analysis, the maximum peak of calcium-dependent signaling for cued licks was significantly different from signaling of uncued licks [p<0.0001], CS+ onset [p = 0.003], and CS+ offset [p = 0.0004]. Thus, BNST GABA neurons show elevated neuronal activity during reward consumption when a few paired backwards CS+ is presented.

**Figure 3.**
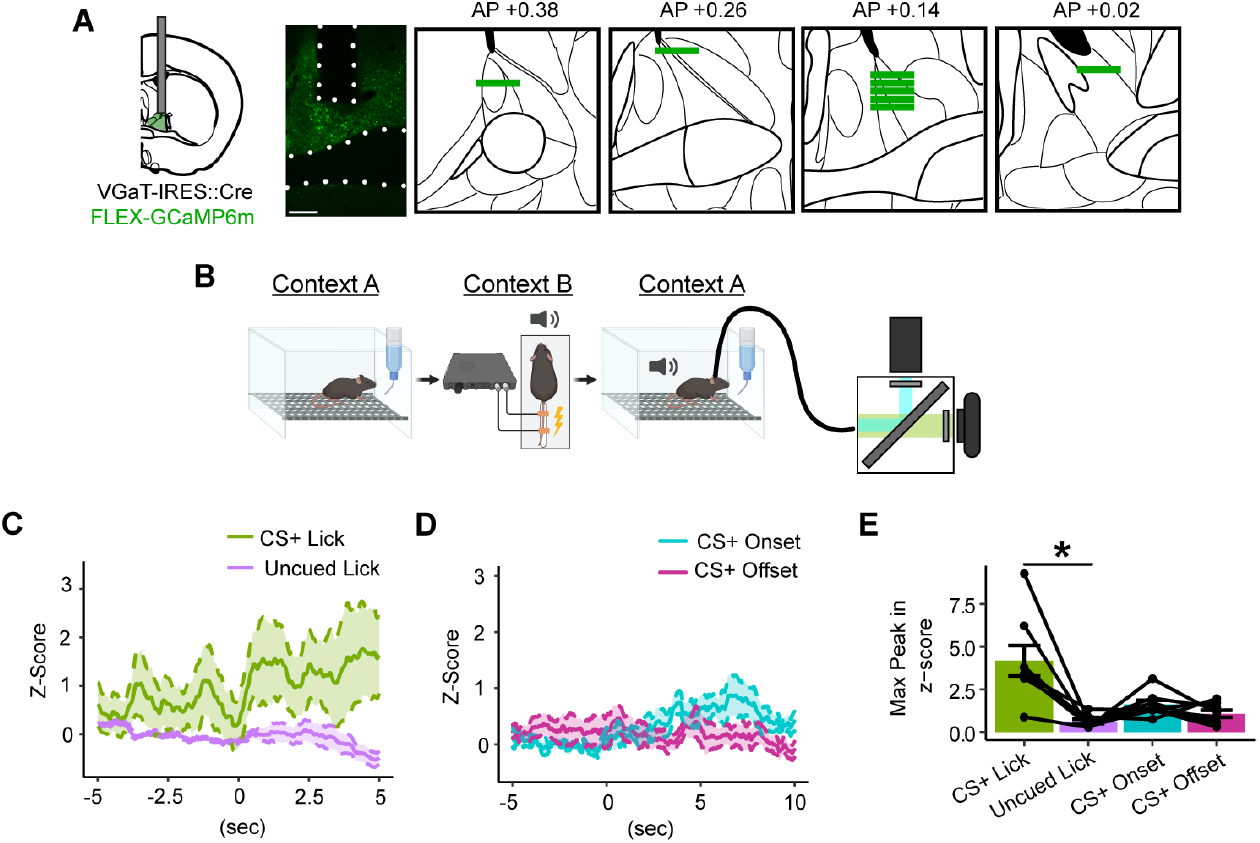
VGaT BNST calcium-dependent signaling during backward conditioned suppression. (A) VGaT-IRES::Cre mice were injected with Cre-dependent GCamP6m. Histology verification of fiber placement are depicted. Scale bar is 200 μm. (B) Illustration of experimental timeline, in which mice were recorded while undergoing the conditioned suppression test. (C) Averaged traces of VGaT calcium-dependent signaling in the BNST in response to CS+ licks and intertrial licks. (D) Averaged traces of VGaT calcium-dependent signaling in the BNST in response to CS+ onset and offset. (E) Maximum peaks observed in response to CS+ licks, intertrial licks, CS+ onset, and CS+ offset. * p < 0.05

### Activation of VGaT BNST signaling abolished backward conditioned suppression

The systemic administration of diazepam, which acts as a positive allosteric modulator of GABA on GABA-A receptors, can have broad effects. However, a subset of BNST GABA neurons are highly activated by diazepam (D. Lu et al., 2024). In order to determine the role of BNST GABA neurons in backwards conditioned suppression, VGaT-IRES::Cre mice were injected in BNST with AAVs encoding Cre-dependent mCherry, hM3Dq-mCherry, or hM4Di-mCherry (**Figure 4A**). After learning to consume sucrose in context A and receiving few backwards pairings of CS+ to tailshock in context B, mice were I.P. administered a behaviorally-subthreshold dose of clozapine to activate the designer receptors (0.1 mg/kg clozapine (Gomez et al., 2017)) 30 minutes prior to undergoing backward CS+ presentations in context A to test conditioned suppression (**Figure 4B**). Prior to conditioning and the conditioned suppression test, there were no group differences in the total number of licks (**Figure 4C**). During the conditioned suppression test, there was a significant effect of DREADDs on CS+ licks [F(2,19) = 3.33, p = 0.05; mCherry = 95.0±14.7, hM4Di = 116.3±24.7, hM3Dq = 182.0±32.2], and post-hoc analysis showed an enhanced number of cued licks for the hM3Dq group compared with mCherry controls [p = 0.05] (**Figure 4D**). Thus, the activation of BNST GABA neurons, not their inhibition, abolished the backward conditioned suppression effect.

**Figure 4.**
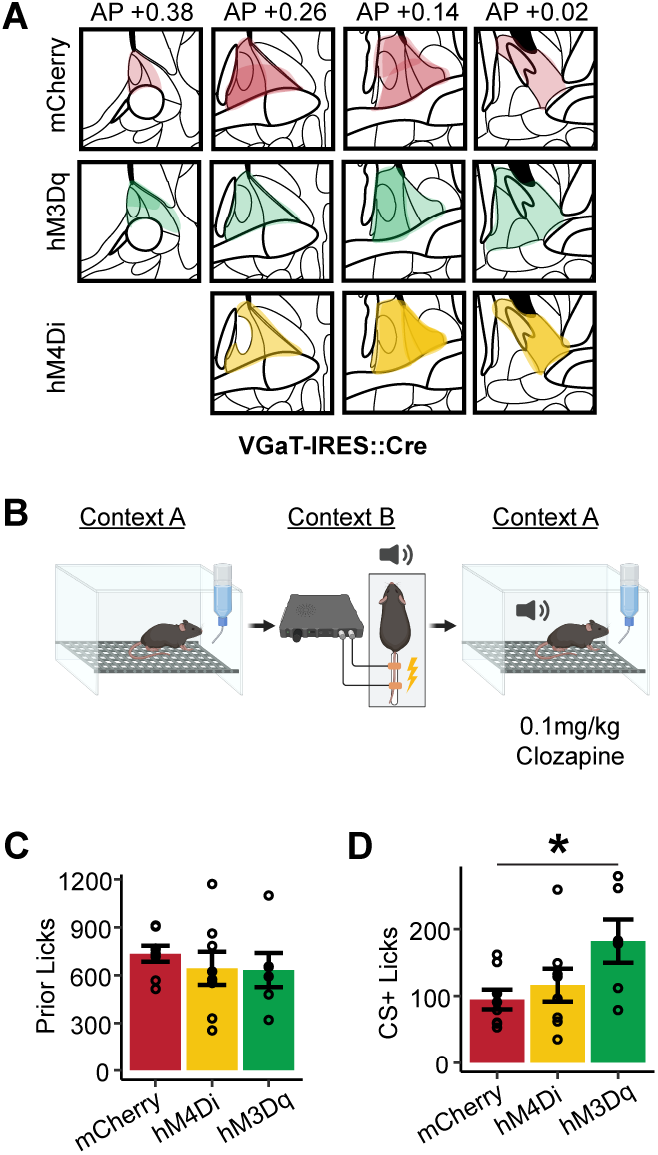
DREADDs activation and inhibition on backward conditioned suppression. (A) VGaT-IRES::Cre mice were injected with Cre-dependent mCherry, hM3Dq, or hM4Di. (B) Illustration of experimental timeline, in which mice were I.P. administered 0.1mg/kg clozapine 30 minutes prior to undergoing the conditioned suppression test. (C) Number of licks prior to conditioning. (D) Licks during CS+ the conditioned suppression test. * p < 0.05

## DISCUSSION

How the history of backwards and forwards conditioning affects conditioned suppression is unknown. We find that few or many forward CS+ pairings and few backward CS+ pairings produced conditioned suppression while many backwards CS+ pairings does not produce conditioned suppression. With forward paired cues, conditioned suppression scales non-linearly with shock probability (Ray et al., 2020; Wright & McDannald, 2019). Forward paired cues with high shock probability results in total conditioned suppression while less-than-chance shock probability (50%) has less conditioned suppression but more than a zero-probability shock CS+. For backwards paired cues, based on the sometimes-opponent process theory, the number of trials is critical to the transition of a backward CS+ from conditioned excitor to conditioned inhibitor (Wagner, 1981). In other words, as subjects learned that a backwards cue indicates shock termination or that the probability of shock is zero following the cue, the backward CS+ transitioned from conditioned excitor to conditioned inhibitor. During few pairings of the backwards CS+ with shock, backward conditioning has been posited to be a conditioned excitor due to the unpredictability of the cue when compared to forward conditioning, which has high predictive value (Goode et al., 2019; Ressler et al., 2020). Alternatively, additional factors influencing the dichotomous conditioned responding effects of backward CS+ may be explained by its association with the context. A few backward CS+ pairings may become associated with the context itself while many backward CS+ pairings may become associated with the intertrial interval that follows and thus becomes a “safety” cue that signals the absence of the US (Chang et al., 2003; Ewing et al., 1985). Indeed, with trace conditioning, the longer the interstimulus interval between the CS+ and US, the more that the CS+ is associated with the context, as demonstrated through contextual fear conditioning (Marlin, 1981). When extinction to the training context was implemented, backward conditioned responding decreased (Chang et al., 2004). Additionally, backward conditioning, regardless of sex, results in higher contextual fear expression than forward conditioning (Olivera-Pasilio & Dabrowska, 2023). An additional factor that may influence the transition of backwards cues from conditioned excitation to conditioned inhibition is the difference in conditioning time between few pairings and many pairings (Gallistel & Gibbon, 2000; Harris & Bouton, 2020). However, this was not the case for forward paired cues that resulted in similar levels of conditioned suppression with few or many pairings. Nevertheless, because few pairings of backwards conditioned stimuli results in either ambiguous or “contextual” relationships with threats, while many pairings result in less ambiguous conditioned inhibition or “safety”, threat-related backwards cues exist at a rare intersection of anxiogenesis and anxiolysis.

Diazapam, a benzodiazepine that is also a treatment option for anxiety disorders (WHO Model List of Essential Medicines - 23rd list, 2023), was used to determine the role of GABAergic signaling in forward and backward CS+ conditioned suppression. Diazepam prevented conditioned suppression to a few backward pairings and not to a few forward pairings. These results suggest that the conditioned suppression behavior produced by a backward CS+ is an anxiety-like behavior that can be rescued with diazepam. We also found that diazepam did not impact freezing behavior, a defensive fear response (Blanchard et al., 2001; Killcross et al., 1997) that was significantly increased in the mice conditioned to few forward CS+ but not in backwards conditioned mice. Nor did diazepam impact distance traveled during the CS+ for forward and backward conditioning. We interpret these results such that: (1) though both few forward CS+ and few backward CS+ cause conditioned suppression, forward cues result in fear-evoked suppression while backward cues result in anxiogenesis-evoked suppression and (2) the influence of diazepam on backward conditioned suppression was based on its ability to increase reward-seeking rather than locomotor activity. At the dose used (0.5mg/kg), diazepam is effective at reducing anxiety-like behavior on the elevated plus maze in both male and female mice (Mehrhoff et al., 2023), supporting that backwards cues may evoke anxiety-like behaviors over forwards cues. In our experiments, diazepam did not change total distance traveled, average speed, and time spent oriented towards the sucrose. Thus, the dose of diazepam used was non-sedative, yet significantly increased reward-seeking during the CS+ for mice conditioned to the backward CS+ and not for mice conditioned to the forward CS+. Given that backwards and forwards cues resulted in differential sensitivity to diazepam and different behaviors that underlie their conditioned suppression, our results support the idea that conditioned freezing and conditioned suppression are separable behaviors that are mediated by different neural circuits (Amorapanth et al., 1999; Blanchard et al., 2001; Killcross et al., 1997; McDannald, 2010; McDannald & Galarce, 2011).

The bed nucleus of the stria terminalis (BNST) is an extended amygdala brain region that is linked to threat uncertainty and anxiety in humans (Awasthi et al., 2020; Brinkmann et al., 2017; Buff et al., 2017; Feola et al., 2023; Mobbs et al., 2010; Naaz et al., 2019; Petranu et al., 2024), as well as in animal models (Davis et al., 2010; Glover et al., 2020; Ly et al., 2023; Sajdyk et al., 2008; Waddell et al., 2006). It is also known that the alpha2 subunit of the GABA-A receptor, which mediates the anxiolytic action of benzodiazepines, is more localized in the BNST than the alpha1 subunit, which mediates the sedative properties of benzodiazepines (Kaufmann et al., 2003). Here, BNST VGaT neurons signaled to sucrose consumption during the backward CS+ but not sucrose consumption outside of the CS+. There was also no change in calcium-dependent signaling of BNST VGaT neurons to backward CS+ onset and offset. The lack of responsiveness to the backwards cue itself was not entirely unexpected because the BNST is largely unresponsive to aversion-predictive forward CS+ (Duvarci et al., 2009; Goode et al., 2019; Haufler et al., 2013; Sullivan et al., 2004). While others have found that backward cues selectively increase c-Fos expression in the BNST (Goode et al., 2019), we find that BNST VGaT calcium-dependent signaling during backward CS+ requires some level of related actions. Further, while pan-neuronal BNST signals to feeding has been previously observed (de Araujo Salgado et al., 2023; Douglass et al., 2023; Jaramillo et al., 2020; Jennings, Rizzi, et al., 2013; Jia et al., 2022; Luskin et al., 2021; Zhang et al., 2023), the subset of BNST neurons we recorded were more sensitive to the interaction of backward cue presentation and consumption.

Systemic administration of diazepam activates the BNST (Dongye Lu et al., 2024), but diazepam can have broad effects that are not exclusive to the activation of the BNST. Therefore, we tested the sufficiency of activating the BNST in abolishing backward conditioned suppression. Using DREADDs, we found that VGaT BNST activation, but not its inhibition, prevented backward conditioned suppression. These results conflict with previous findings that BNST activation produces anxiogenic behavior (Mazzone et al., 2018; Williford et al., 2023; Yamauchi et al., 2018). Methodological differences between these and the current experiment may explain this difference. For example, the previously noted studies that assessed anxiety-like behavior used classic self-guided novel contexts that would normally be anxiogenic to rodents. Using these classical tasks, BNST activation results in anxiogenic behavior. Here, the backward CS+ that is associated with shock is presented in conflict to the context, which is rewarding and familiar. Under these circumstances, BNST activation prevented anxiety-like behavior observed during backward CS+ presentation by increasing reward intake that would normally be observed in response to a conditioned inhibitor. Taken together with prior literature, our results suggest a nuanced role of BNST in anxiety and in conflicts of reward and aversive processing. One limitation of the current experiments was that we assessed VGaT BNST-specific neurons while differential behaviors may result from the activation of molecularly-distinct subsets (Jennings, Sparta, et al., 2013) or subregions (Kim et al., 2013) of BNST VGaT neurons.

## ACKNOWLEDGEMENTS

This research was supported by the Webb-Waring Biomedical Research Award from the Boettcher Foundation (DHR), Institute for Cannabis Research (DHR), National Institute on Drug Abuse DA047443 (DHR), MH130576 (DHR), the University of Colorado ABNexus (DHR), and the National Institute on Mental Health MH13222 (AL). AL and DHR contributed to experimental design, data interpretation, and manuscript writing. AL, DHR, HH, EDP, JP, MD, and LM contributed to data collection. All authors edited and reviewed the manuscript for publication.

